# Transposable element over-accumulation in autopolyploids results from relaxed purifying selection and provides variants for rapid local adaptation

**DOI:** 10.1101/686584

**Authors:** P. Baduel, L. Quadrana, B. Hunter, K. Bomblies, V. Colot

## Abstract

Polyploidization is frequently associated with increased transposable element (TE) content. However, what drives TE dynamics following whole genome duplication (WGD) and the evolutionary implications remain unclear. Here, we leveraged whole-genome resequencing data of ∼300 individual *Arabidopsis arenosa* plants, a well characterized natural diploid-autotetraploid species, to address these questions. Based on 43,176 polymorphic TE insertions we detected in these genomes, we demonstrate that relaxed purifying selection rather than transposition bursts is the main driver of TE over-accumulation after WGD. Furthermore, the increased pool of TE insertions in tetraploids is especially enriched within or near abiotic and biotic stress response genes. Notably, we uncovered one such insertion in a major flowering-time repressor gene and found that the resulting allele is specific to the rapid-cycling tetraploid lineage that colonized mainland railways. Together, our findings indicate that tetrasomy by itself leads to an enhanced tolerance to accumulating genic TE variants, some of which can potentially contribute to local adaptation.

## Introduction

Eukaryotic genomes display remarkable variation in ploidy as well as in size. This is particularly true of plant genomes, the evolution of which has been punctuated by numerous whole genome duplication (WGD) events^1–3^. Moreover, polyploidization is often associated with an increase in transposable element (TE) content, which further exacerbates the inflation of genome size^3,4^. The frequent association between polyploidization and higher TE content has long been seen as evidence for the “genome shock” hypothesis^5^, where genomic instabilities associated with genome doubling could trigger transposition bursts. Yet, the possibility of such bursts has been studied primarily in allopolyploids, where the effects of WGD and hybridization are confounded (e.g.^6,7^). Furthermore, gene redundancy is expected to reduce the selective pressure exerted on recessive deleterious mutations^8^ especially in autopolyploids where all homologous chromosomes segregate randomly (polysomic inheritance). As a result, the loss of function mutations typically caused by TE insertions could readily over-accumulate in polyploids, even in the absence of transposition bursts.

These two non-mutually exclusive scenarios are still presented indiscriminately (e.g.^9^) because of a lack of experimental support of one or the other^10,11^. Yet, the implications for the evolution of polyploids differ substantially under the two models: transposition bursts resulting from the WGD event are expected to impact the fitness of neo-polyploids in the first few generations after they are formed^12,13^, while the effects of relaxed purifying selection would be perceived many generations downstream, as a result of the progressive accumulation of new TE insertions. Distinguishing between these two possibilities is therefore critical for our understanding of the evolutionary implications of ploidy-associated TE variation.

Here, we set out to assess what drives TE dynamics in *Arabidopsis arenosa*, a model system for studying WGD independently of the confounding effects of hybridization. This plant species occurs in both diploid and autotetraploid forms across Central and Northern Europe. The autotetraploid originated in a single recent WGD event^14^ (∼60kyr ago) and has rapidly expanded, thus occupying a broader ecological range than the diploid progenitor^15^. Notably, and as frequently reported for polyploids^16,17^, one *A. arenosa* tetraploid lineage successfully invaded a novel ruderal habitat, where they adopted a weedy life-style life-cycle not shared by diploids or non-ruderal tetraploid *A. arenosa*^*18,19*^.

Using the most comprehensive diploid-autopolyploid dataset to date^20^, we characterized TE dynamics across diploid and tetraploid *A. arenosa* genomes. First, we assessed how the relaxation of purifying selection impacted TE dynamics in autotetraploids. Second, we investigated whether a transposition burst was associated with the WGD event that all *A. arenosa* tetraploids trace back to. Finally, we asked whether and how TE dynamics in autotetraploids might contribute to their adaptive potential.

## Results

### Non-reference TE landscape across 286 diploid and tetraploid *A. arenosa* genomes

Genome resequencing data are available for 105 diploid and 181 tetraploid individuals of *A. arenosa*, corresponding to 15 and 23 distinct populations covering the entire range of the species, respectively^20^. In the absence of a reference *A. arenosa* genome sequence, we used that of the closely related *Arabidopsis lyrata*^*2121*^ as was done in previous SNP-based population genomic studies of *A. arenosa*^*14,20,22*^.

We assessed recent TE activity in *A. arenosa* using the SPLITREADER pipeline^23^ (see Methods), which enabled us to identify 43,176 non-reference TE insertions from split and discordant reads in the 286 re-sequenced *A. arenosa* genomes aligned on the *A. lyrata* genome. These TE insertions correspond to three class I (*LINE, Copia*, and *Gypsy* retrotransposons) and five class II (*Mariner, MuDR, hAT, Harbinger*, and *CACTA* DNA transposons) superfamilies, in proportions similar to those found in *A. lyrata*^*24*^ (Fig. 1A). However, unlike annotated TE sequences of the *A. lyrata* reference genome, which are heavily enriched towards pericentromeric regions, the non-reference TE insertions we detected are homogeneously distributed along chromosomes with no obvious pericentromeric bias (Fig. 1B).

**Figure 1.**
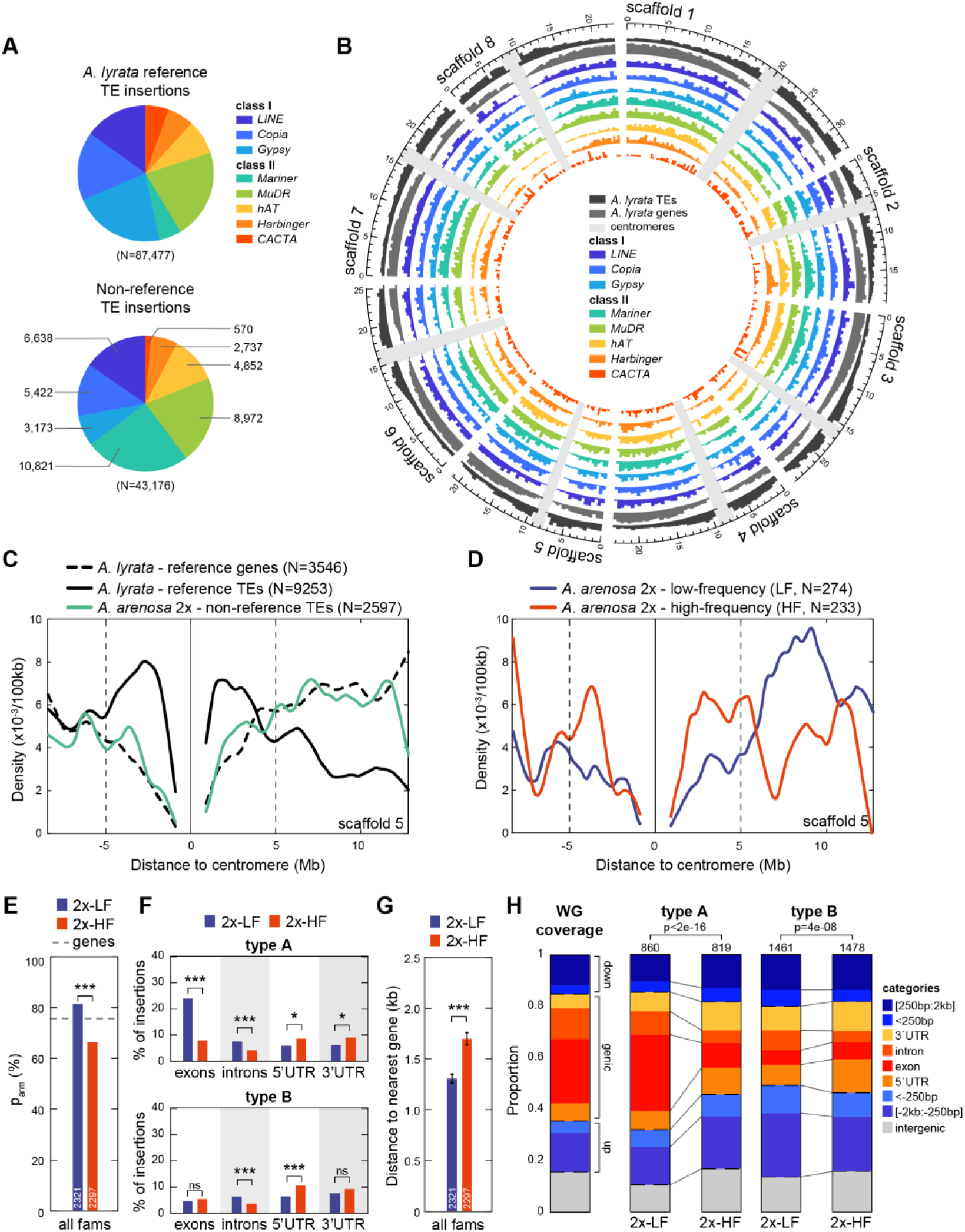
Natural selection shapes the TE landscape of *A. arenosa* diploids. (A) Distribution of reference (upper chart) and non-reference (lower chart) TE insertions identified by SPLITREADER across the 8 class I & II TE superfamilies. (B) Chromosomal distributions of reference genes and TEs and of non-reference TE insertions by TE superfamily across the 8 scaffolds of the *A. lyrata* reference genome. (C) Density per 100kb of reference genes and TEs and of non-reference diploid TE insertions across scaffold 5. (D) Density per 100kb across scaffold 5 of low- and high-frequency non-reference TE insertions in diploids. (E) Fraction, p_arm_, within chromosome arms (>5Mb away from centromeres) of low- and high-frequency non-reference TE insertions in diploids. (F) Fraction of low- and high-frequency non-reference TE insertions in diploids overlapping exons, introns, 5’ or 3’ UTRs for type A (upper panel) and type B (lower panel) superfamilies. (G) Distance to nearest gene (kb) of low- and high-frequency non-reference non-genic TE insertions in diploids. (H) Distribution of low- and high-frequency non-reference TE insertions in diploids across categories of insertions for type A and type B superfamilies compared to reference genome annotations. (p<0.001: ***; p<0.01: **; p<0.05: *; p≥0.05: ns)

### Purifying selection shapes the TE landscape of *A. arenosa* diploids

To more precisely characterize TE localization and dynamics separately we started with the 105 diploid (i.e. 2x) individuals. The density of non-reference TE insertions is higher on the chromosome arms than within 5Mb of the centromeres and follows that of the genes rather than the TEs annotated in the *A. lyrata* reference genome (Fig. 1C). In order to test whether this higher density of non-reference TE insertions within chromosome arms reflects recent TE activity, we compared the distribution along chromosomes of those TEs at low- and high-frequency (2x-LF vs 2x-HF, see Methods). This parameter is a proxy for relative age when selection is not a factor, because recent insertions are expected to be at lower frequency than older ones. Across all TE superfamilies, we observed a clear and consistent decrease (−13.5%, p<0.001) in the density and proportion (parm) of HF insertions within chromosome arms compared to LF insertions (Fig. 1D-E). This distribution shift was also associated with a net deficit of HF TE insertions within or near (<250bp) genes (Fig. 1F-G); HF genic TE insertions are also strongly depleted from exons and introns (Fig. 1F). These observations are consistent with purifying selection acting to progressively filter out TE insertions from chromosome arms due to their generally deleterious effects on genes, as reported in *A. thaliana*^*23*^. In keeping with a potentially mutagenic cost of transposons, in our previously published RNAseq datasets available for a subset of nine populations^19,20^, we found a significant transcriptomic impact of non-reference TE insertions on gene expression when these are located within or near (<250bp) genes but not when further away (Fig. S1, see Methods).

Further analysis revealed that the deficit of HF TE insertions within or near genes is most pronounced for the *Copia, Gypsy, CACTA*, and *hAT* TE superfamilies, which show an apparent insertion preference for genes, especially exons, and are hereafter called ‘type A’ (Fig. H-S2). Specifically, the proportion of non-reference type A insertions that lie within exons and introns are much reduced (−20.0% and −4.2%, respectively) at high frequency (Fig. 1F), consistent with them having the most detrimental effects and therefore being the most rapidly purged by natural selection.

In contrast, non-reference TE insertions from the other TE superfamilies (*LINE, Mariner, MuDR*, and *Harbinger*), thereafter called ‘type B’, are underrepresented within exons whether at low or high-frequency (Fig. F-H-S2). It is not clear whether this observation reflects insertion preferences away from exons or stronger and more dominant deleterious effects of type B exonic insertions, which would thus be rapidly purged and therefore not detected even at low-frequency.

The deficit of HF TE insertions within exons (type A) or introns (types A&B) is not compensated uniformly across other categories. Rather, an increased proportion of HF TE insertions was most pronounced in the 5’ or 3’ UTRs of genes for both types of TE superfamilies (+4.5% and +2.6% respectively, Fig. 1F-S2). This observation suggests that insertions located within these two particular genic compartments may more frequently be under positive rather than purifying selection.

### Increased non-reference TE content in autotetraploids results from relaxed purifying selection, not transposition bursts

In contrast to diploids, autotetraploids (i.e. 4x) show a marked increase in the proportion of non-reference type A TE superfamily insertions within exons (Fig. 2A-B). In addition, non-reference type A insertions are slightly more frequent within chromosome arms (p<0.05, Fig. 2C) and closer to genes (0.10kb, p<0.001, Fig. 2D) in the autotetraploids relative to diploids. The proportion of non-reference type A insertions within UTRs are mostly unaffected by ploidy (Fig. 2A). In contrast to non-reference type A insertions, those of type B showed few differences between ploidies, except for a small increase in their proportion within UTRs in the tetraploids (Fig. 2E), the significance of which is unclear. This last observation notwithstanding, our findings are consistent with purifying selection being significantly relaxed on genic TE insertions in the autotetraploids.

**Figure 2.**
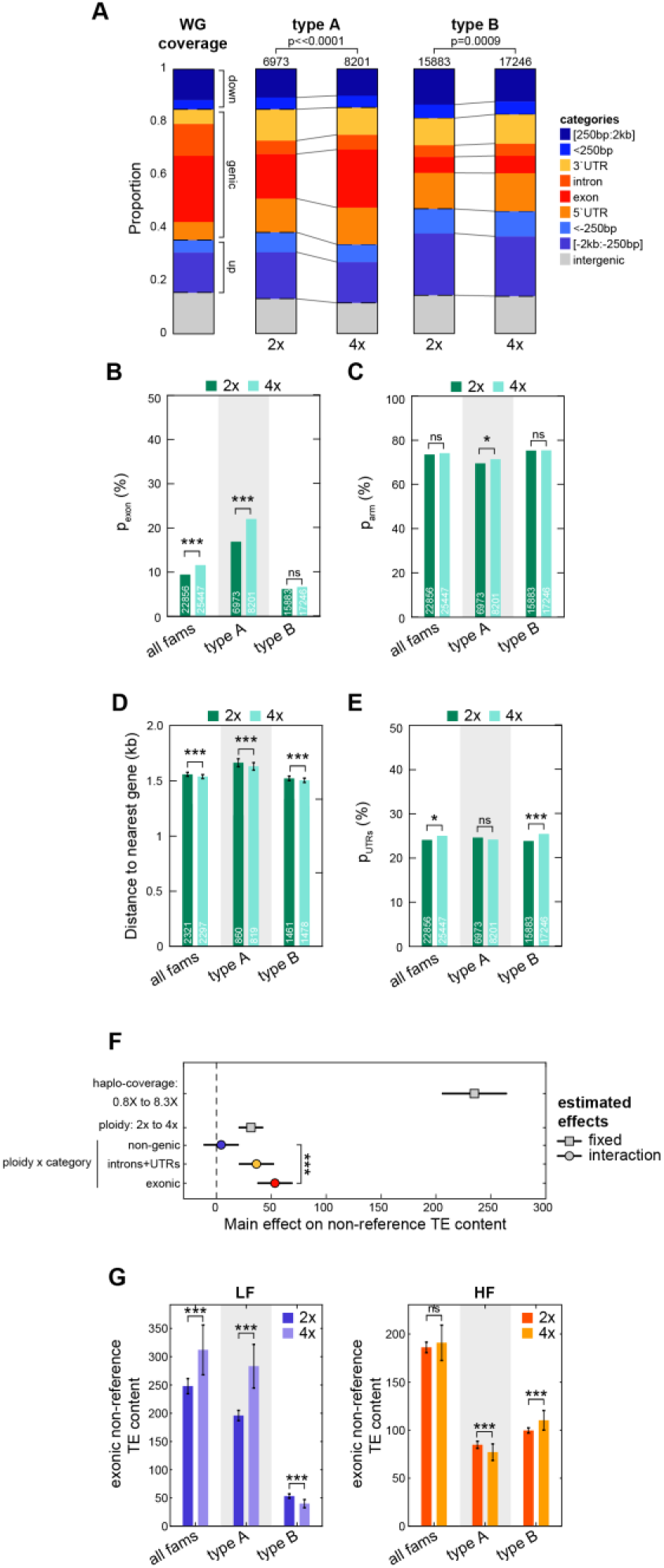
Increased exonic TE load from relaxed purifying selection in autotetraploids. (A) Distribution of non-reference TE insertions across categories of insertions for type A and type B superfamilies in diploids and tetraploids compared to reference genome annotations. (B) Fraction, p_exon_, of non-reference TE insertions overlapping exons for all, type A, and type B superfamilies in diploids and tetraploids. (C) Fraction, p_arm_, within chromosome arms (>5Mb away from centromeres) of non-reference TE insertions for all, type A, and type B superfamilies in diploids and tetraploids. (D) Distance to nearest gene of non-reference non-genic TE insertions for all, type A, and type B superfamilies in diploids and tetraploids. (E) Fraction, p_UTRs_, of non-reference TE insertions overlapping UTRs (5’ and 3’) for all superfamilies, type A, and type B TE superfamilies in diploids and tetraploids. (F) Estimated MLM effects and interaction terms of haplo-coverage, ploidy, and category of insertions (non-genic, introns and UTRs, or exonic) on non-reference TE content. (G) Number of non-reference TE insertions within exons carried by 100 individuals for all, type A, and type B superfamilies in diploids and tetraploids at low-frequency (LF, left panel) and high-frequency (HF, right panel). (p<0.001: ***; p<0.01: **; p<0.05: *; p≥0.05: ns)

Consistent with this scenario, autotetraploids tend to harbor a higher number of non-reference TE insertions per haploid genome than their diploid counterparts (see Methods, Fig. 2F, Table S1) and this increase is predominantly contributed by exonic insertions. Furthermore, tetraploids on average carry∼30% more low-frequency TE insertions (mostly type A) within exons than diploids (see Methods, Fig. 2G), but the number of high-frequency TE insertions is similar between diploids and autotetraploids (Fig. 2G). Given that autotetraploids arose relatively recently ^14^ and are therefore unlikely to have reached their mutational balance equilibrium ^20^, the relaxation of purifying selection should be mostly visible over low-frequency variants, as these tend to be young and not shared with diploids.

According to the genome-shock hypothesis, the increase in non-reference TE content observed in the autotetraploids could also be the footprint of a general transposition burst at the time of the WGD event. Under this scenario, tetraploids compared to diploids should exhibit an excess of high-frequency insertions, especially when non-genic, as these are the least exposed to purifying selection (Fig. 1H). This is clearly not the case, as we observed an excess of such insertions in the tetraploids at low-frequency only (see Methods, Fig. 3A-B, Table S2). Nonetheless, 35% of high-frequency non-genic insertions in the autotetraploids are not present in the diploids (Fig. S3), suggesting that they could have originated from TE family-specific transposition bursts. To address this, we compared the proportion of high-frequency non-genic insertions between tetraploids and diploids for the 201 TE families with enough non-reference copies (>10) in each of the two ploidy groups and found that no TE family shows a significant excess (χ^2^, p<0.05) of high-frequency non-genic insertions in the autotetraploids, with one possible exception (*ALLINE1_2*; p=0.036, Fig. 3C).

**Figure 3.**
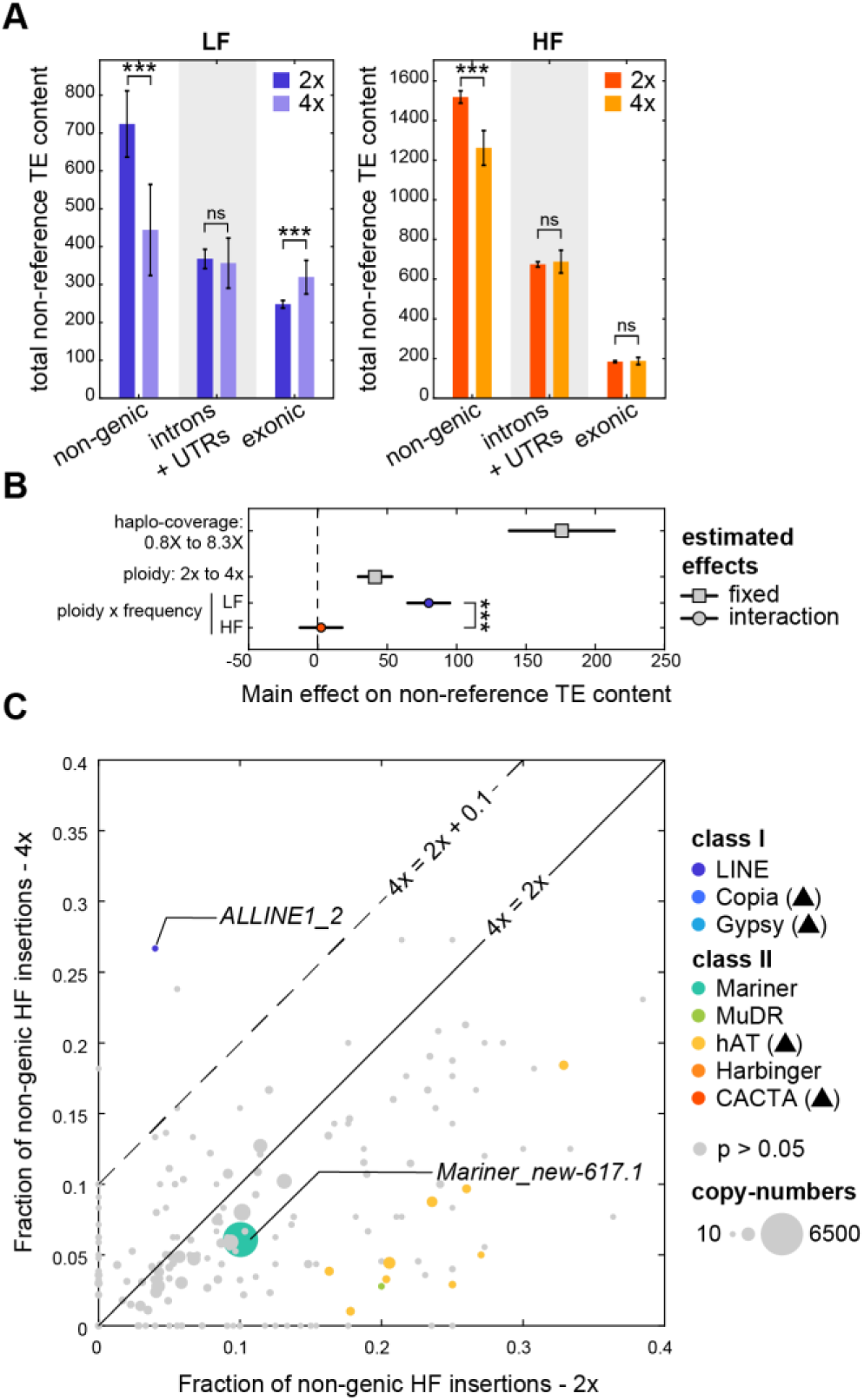
In the absence of transposition burst hallmarks, relaxation of purifying selection explains increased non-reference TE content in autotetraploids. (A) Number of non-reference TE insertions carried by 100 individuals by category (non-genic, introns and UTRs, exonic) in diploids and tetraploids at low-frequency (LF, left panel) and high-frequency (HF, right panel). (BF (C) Fraction of non-genic insertions at high-frequency (HF) in tetraploids versus diploids by TE-family. TE-families with χ^2^ p-values < 0.05 are colored.

Collectively, our data indicate that the increased non-reference TE content in autotetraploids results mainly if not exclusively from the progressive accumulation of insertions within genes, especially exons, thanks to relaxed purifying selection, rather than from any appreciable transposition burst.

### Increased *Copia* accumulation in tetraploids provides variants for local adaptation

Given the association of polyploidy with colonization potential^25–27^, we asked whether the increased non-reference TE content within or near genes could contribute to local adaptation of autotetraploids. To this end, we first identified clade-specific TE insertions for the two ploidy groups (See Methods, Fig. 4A) and found a significant enrichment in tetraploids compared to diploids within or near (<250bp) genes, which is most pronounced for local high-frequency type A insertions (Fig. 4B). This finding suggests therefore that type A insertions within or near genes are commonly under local positive selection in tetraploids, unlike those of type B (Fig. 4B). Further analysis of these local high-frequency type A insertions indicated that they are specifically enriched for *Copia* LTR-retrotransposons in tetraploids (Fig. 4C). Consistent with observations in *A. thaliana* and other plant species^28^, this TE superfamily shows in both ploidies a marked insertion preference towards genes responsive to biotic stimuli (Fig. S4), a class of genes that are typically under diversifying selection^29,30^. Moreover, *Copia* insertions within or near genes of this category are significantly enriched at local high-frequency only in tetraploids (Fig. 4D) and are more likely to reach high frequencies (3 or more individuals within a clade, HF3) than all other insertions (Fig. 4E). Collectively these findings indicate that *Copia* insertions within stimulus response genes are more frequently under local positive selection specifically in tetraploids.

**Figure 4.**
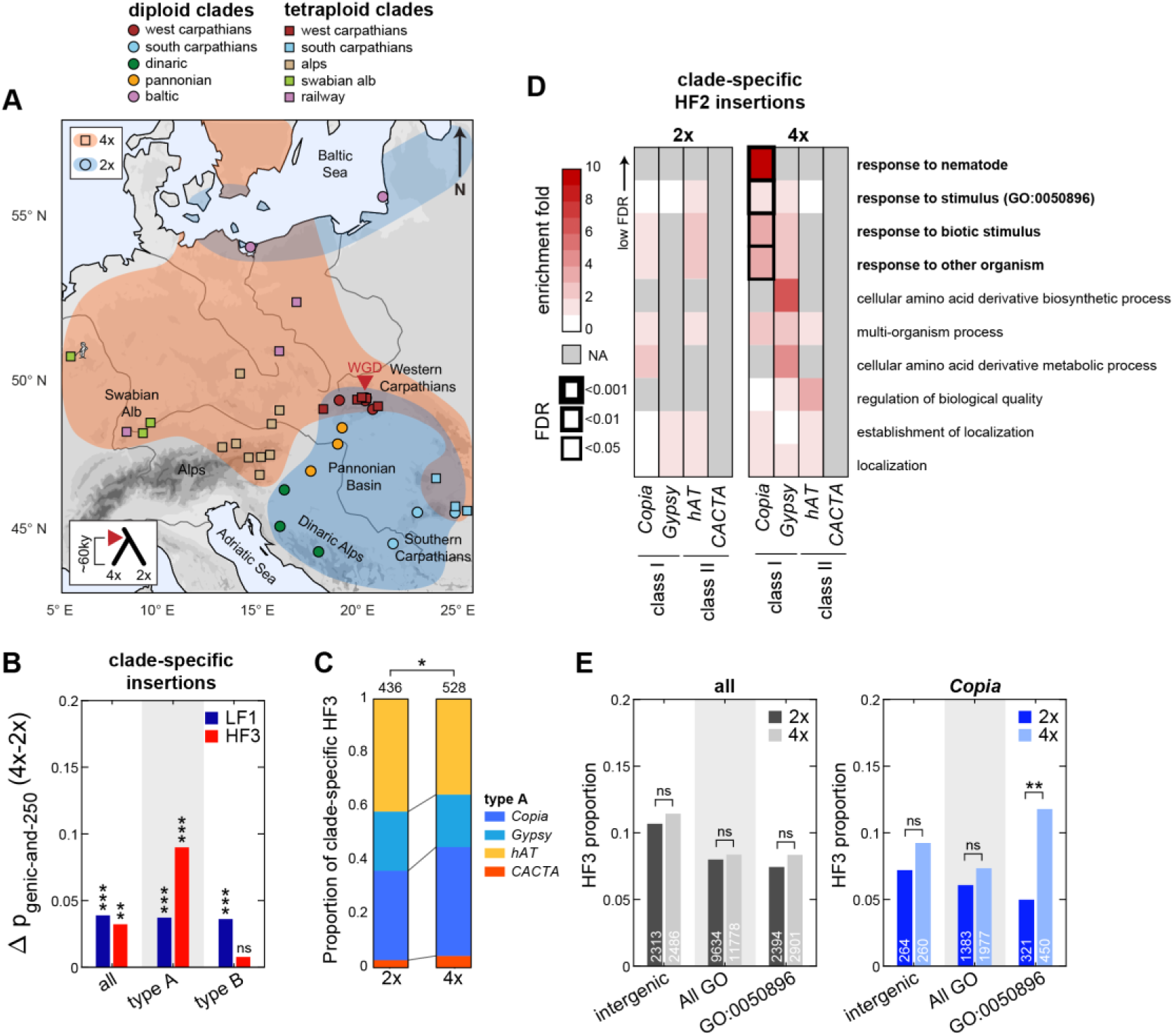
Local positive selection on *Copia* genic insertions in tetraploids. (A) Map of the tetraploid and diploid populations colored by clade. (B) Difference between tetraploids and diploids of the proportion of genic and near-genic (<250bp) insertions present in only 1 clade (clade-specific) and carried by no more than 1 individual (LF) or by 3 or more individuals within the clade (HF) for all, type A, and type B superfamilies. (C) Proportion among clade-specific HF3 type A insertions from each type A superfamily in diploids and tetraploids. (D) GO enrichments in diploids and tetraploids among genes carrying or nearby clade-specific HF2 insertions for type A TE superfamilies. (E) Proportion in diploids and tetraploids of clade-specific HF3 insertions within intergenic regions, within or near genes, or within or near stimulus response genes (GO:0050896) for all superfamilies or only *Copia* insertions. (p<0.001: ***; p<0.01: **; p<0.05: *; p≥0.05: ns)

One clear example of local adaptation of *A. arenosa* tetraploids to a novel habitat is provided by the successful colonization by one lineage of railway ballasts across central Europe. In previous studies, we have shown that this adaptation is associated with a switch to early flowering and a loss of expression of the floral repressor *FLC*^*18,19*^. *FLC* is a known hotspot for *Copia* insertions in *A. thaliana*^*23*^, so we investigated whether railway tetraploids may be carrying such an insertion. Due to local syntenic divergences between *A. arenosa* and *A. lyrata* in the *FLC* region^31^, we could not use our *A. lyrata-*based pipeline to search for the presence of non-reference TE insertions within the 2 full length *A. arenosa FLC* paralogues (*AaFLC1* & *AaFLC2*). Instead, we sequenced three fosmid clones of the *FLC* region that were obtained from an early-flowering railway individual. Comparison to the BAC sequence of the *FLC* region that was obtained previously from a late flowering individual^31,32^, revealed a number of intergenic or intronic structural variants. Notably, two clones contained the same *ATCOPIA78* solo-LTR insertion in the 2^nd^ exon of *AaFLC1* (Fig. 5A), which is the main contributor (>80%) to total *FLC* expression in *A. arenosa*^*18*^. Examination of whole-genome sequencing data of the 286 *A. arenosa* accessions for the presence of this solo-LTR insertion (see Methods) indicated that it is present in the three railway tetraploid populations, but not in any of the 15 diploid populations nor in any of the other 19 tetraploid populations analyzed (Fig. 5B). It was also absent from a hybrid mountain-railway population, which, unlike the other railway tetraploids, is characterized by high *FLC* expression and was shown previously to be early flowering because of a specific allele of *CONSTANS*, another major flowering time regulator^19^. The complete association between the *Copia* insertion and the loss of *FLC* expression provides a strong argument in favor of the potential of *Copia* retrotransposons for generating alleles that can enable rapid adaptation to novel habitats.

**Figure 5.**
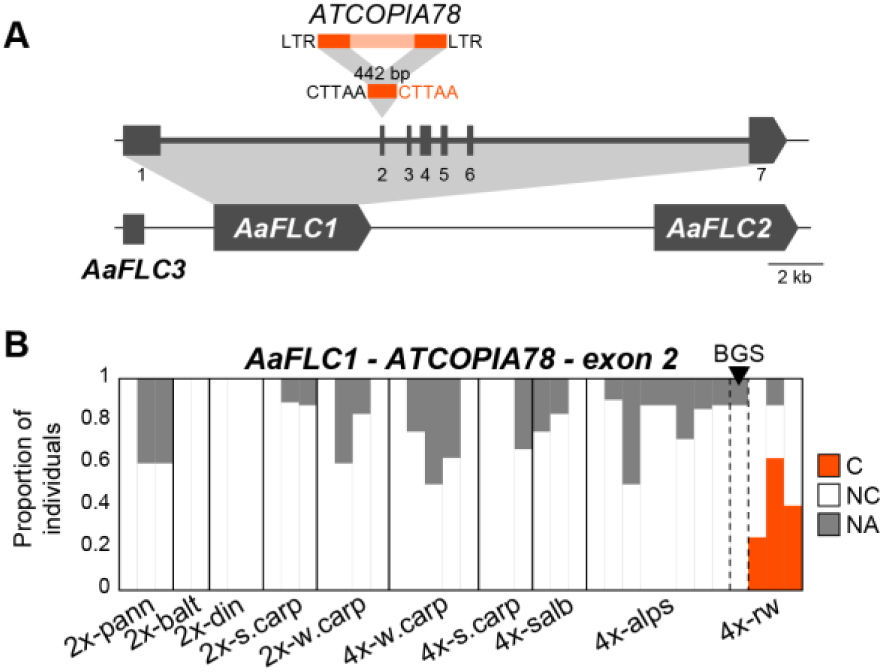
Railway-specific *ATCOPIA78* solo-LTR exonic insertion within *FLC*. (A) Map of *FLC* region with 442bp-long *ATCOPIA78* solo-LTR insertion in 2^nd^ exon of *AaFLC1* identified by fosmid sequencing. (B) Proportion of individuals across populations by clade carrying the *ATCOPIA78* solo-LTR insertion. Hybrid mountain-railway population BGS is indicated with black triangle.

The ability of tetraploids to absorb locally-adaptive variants through introgression with local diploids has been often argued to favor their potential for rapid adaptation^20,33^. At the TE level, we could confirm the strong admixture between southern Carpathians diploids and tetraploids (Fig. S5A). Yet, even in this case, diploid TE variants represented only a minor fraction of TE insertions at local high-frequency within the south Carpathian tetraploids (Fig. S5B) and particularly so for *Copia* which are almost exclusively locally-sourced within the tetraploids.

## Discussion

Here, we performed a comprehensive analysis of TE dynamics in a natural diploid-autopolyploid species, which enabled us to assess the consequences of WGD without the confounding effect of hybridization. Our results indicate a major over-accumulation of TEs specifically within or near genes in the autotetraploids compared to their diploids counterparts, which supports the hypothesis that polysomy shelters TE insertions from selection. Conversely, we found no evidence of transposition bursts, neither genome-wide nor family-specific that could have been associated with the ancestral WGD event, which suggests that genome duplication alone, when not coupled with hybridization as in allopolyploids, is insufficient to cause a severe “genomic stress”. Finally, we present data supporting a role in local adaptation of the genic TE variants accumulating in tetraploids, including within a major adaptive locus.

Our conclusion that purifying selection in autotetraploids is relaxed confirms previous work based on single nucleotide polymorphisms (SNPs)^20^. However, the resulting increase in genetic load remained subtle, which is expected given the young age of *A. arenosa* tetraploids^14^ and the small fitness effects of most point mutations. In contrast, the impact of relaxing purifying selection is magnified for TEs which often engender major-effect alleles. Over longer term evolution, we expect this fundamental difference to be further exacerbated in a chain reaction-like manner since transposition rates may increase as more active copies are tolerated in tetraploid genomes, as has been observed in *A. thaliana* TE-mutation-accumulation lines^28^.

The progressive over-accumulation of genic TE variants seen as a result from relaxed purifying selection could provide raw material for local adaptation of some tetraploid populations. Indeed, we observed that *Copia* insertions within or near stimulus response genes appear to be under local positive selection specifically in tetraploids. In contrast, our results show that interploidy admixture with local diploid populations contribute very little potentially adaptive *Copia* variants. The successful invasion of railway ballasts by *A. arenosa* tetraploids has been particularly intriguing as it represents an example of the increased invasion potential often associated with polyploidy^25–27,34^ though it seems at least in part to rely on diploid alleles imported through admixture^19^. Here, however, we found that a tetraploid-specific *Copia* exonic insertion within *AaFLC1* was both unique to and common in all the rapid-cycling railway populations, with the exception of one hybrid population where early-flowering was not associated with a loss of *FLC* expression^19^. Further work is needed to characterize the case by case phenotypic impact of local TE insertions in particular at the *FLC* locus.

While TEs may provide adaptive opportunities, they can also have negative impacts. In addition to directly producing deleterious mutations, TEs are major contributors of spurious recombination events^9^, a key step in the “polyploid drop”^35^ that has historically almost inevitably followed WGD events^3,36^. Thus TE accumulation in autopolyploids may provide transient windows of adaptive opportunity but could eventually also lead to their evolutionary demise or at least their return to a diploid state.

In summary, our study sheds new light on the dynamic interactions between ploidy and the TE landscape with major implications for the adaptive potential and evolution of polyploids.

## Materials and Methods

### Data sources

Whole genome sequencing data of *A. arenosa* individuals and paired-end alignments on the *A. lyrata* reference genome^20,21^ were obtained from Monnahan *et al.^20^*.

### Identification of non-reference TE insertions

Split and discordant reads were extracted from individual alignments and mapped on a joint TE library assembled from the annotation of all TEs in the *A. lyrata* reference genome (87,477 TEs^24^). Mapped reads were then soft-clipped and re-mapped to the reference genome to define putative non-reference insertion sites. Putative insertion sites were intersected across individuals to define shared insertion sites supported in at least one individual by a minimum of 3 reads, including at least one upstream and one downstream. Negative coverage, as defined by the minimum read depth over the upstream and downstream boundaries of an insertion site, was then calculated for each individual across the putative insertion sites thus obtained. To limit false-negatives, non-carrier individuals with less than 5 reads negative coverage or more than 100 (>10 times the average read depth) were considered as missing information or NA. Sites with more than 10 NA diploids and 10 NA tetraploids were considered non-informative for further analyses, resulting into 43,176 informative insertion sites.

### Analyses of TE landscapes

Statistical analyses were performed using MATLAB (MathWorks, Inc., Natick, Massachusetts, United States). Allele frequency was calculated at each insertion site over all non-NA individuals for the site. Frequency thresholds for LF and HF insertions were calculated as the 10% and 90% percentile of the diploid frequency spectrum (≤1.2% vs ≥8.3%, respectively).

Densities of TE insertions across chromosomes were calculated by 100kb windows and smoothed using a LOWESS (Locally Weighted Scatterplot Smoothing) regression for plotting (Fig. 1C-D). Categories of TE insertions were based on the RNAseq-improved annotation of the *A. lyrata* reference genome^21,37^. In case of ambiguity the following priority order was given: 3’UTR-5’UTR-exon-intron-<250bp upstream-<250bp downstream. For insertions both <2kb upstream of one gene and <2kb downstream of another gene the attribution to one of the two categories was defined by the closest gene. The representations of these categories in the reference genome (WG coverage in Fig. 1H) were obtained following the same rules and were then normalized by their average coverage calculated in 10kb windows using a high-coverage diploid alignment (47X). Pairwise differences in proportions by category (either LF vs HF or 2× vs 4×) were estimated using a 2×2 chi-2 test. Multiple linear models (MLMs) of individual non-reference TE content (Fig. 2F-3B) were obtained by stepwise multiple linear regressions using as parameters haplo-coverage (average coverage by haploid genome), ploidy, and category of insertion (non-genic, exonic, and other genic) for Fig. 2F, or haplo-coverage, ploidy, and frequency (LF and HF) for Fig. 3B (Tables S1-S2). Fixed effects or interaction terms were added or removed based on the p-value for an F-test of the change in the sum of squared error with or without the term. Average TE content by ploidy was calculated based on 100 subsamples of 100 individuals by ploidy. Error-bars indicate standard deviation observed across 100 samples and statistical differences were obtained by 2-way t-tests (Fig. 2G-3A). Gene Ontology (GO) enrichments were calculated based on genes <250bp away from non-reference TE insertions of interest using agriGO in comparison with *A. thaliana* reference annotation (http://bioinfo.cau.edu.cn/agriGO/).

### RNAseq analysis

RNAseq datasets were obtained from previously published datasets^19,20^ for a subset of six tetraploid and three diploid populations with three individuals each. For each gene, we compared the average expression levels in populations where the nearest TE insertion was detected (carrier, C populations) to populations where the nearest TE insertion was not detected (non-carrier, NC). For genes with TE insertions >2kb away (intergenic) C/NC ratio distribution was compared to randomly picked carriers and non-carriers populations (Fig. S2, Kolmogorov-Smirnov test). For genes with TE insertions <250bp away (near-genic insertions), the distribution of C/NC ratios was compared to both random (KS; p<5e-10) and intergenic (Fig. S2, KS p<5e-3).

### Fosmid libraries

Because the *FLC* region in *A. arenosa* contains 3 *FLC* paralogues (two entire copies *AaFLC1* & *AaFLC2* and one truncated *AaFLC3*), that arose independently from the duplicated *A. lyrata FLC* paralogues^31^, Illumina short-reads align poorly on the FLC locus of *A. lyrata* reference genome impairing the detection of non-reference TE insertions. Instead, we extracted DNA from three-week-old plants from a mainland railway population (TBG) using a large-scale CTAB protocol including treatment with pectinase. We constructed a fosmid library using the Copy Control Fosmid Library Production Kit (Epicentre) and screened it using DIG-labeled (Roche) PCR probes to the center of the *FLC* locus (primers 5’ AGTGTAACTTCAATGGCAGAAAACCCT 3’ and 5’ ATGTGGCGGTAAGCAGAGATGACC 3’). We bar-coded positive clones and sequenced 100bp paired-end reads on an Illumina HiSeq 2000. We aligned *FLC* reads (which performed poorly in *de novo* assembly) to an *A. arenosa* BAC (GenBank accession no. FJ461780) using BWA. We identified insertions and deletions by targeted *de novo* alignment using Velvet^38^. The 442bp-long insertion present in the second exon of *AaFLC1* in two out of three fosmids bore a 94.5% similarity with the LTR sequence of *ATCOPIA78* (RepeatMasker blast) in addition to a 5bp tandem site duplication (TSD), as expected from *Copia* insertions^39^.

We detected carrier individuals for this insertion from whole-genome resequencing data by re-mapping paired-end reads to two versions of an updated BAC sequence^18^ including or not the insertion. We considered as carrier any individual with at least 1 read bridging the insertion extremities by at least 20bp, and as non-carrier when no read bridged insertion extremities and at least 4 reads bridged the TSD.

## Acknowledgements

PB would like to thank Levi Yant for discussions and access to unpublished data, Vincent Castric for communication of unpublished *A. lyrata* TE annotation, and Pirita Paajanen for the high-coverage diploid *A. arenosa* alignment.

Work in the Colot lab is supported by the Investissements d“Avenir ANR-10-LABX-54 MEMO LIFE, 506 ANR-11-IDEX-0001-02 PSL* Research University. L.Q. acknowledges support from the MOMENTUM program of the Centre National de la Recherche Scientifique. PB was the recipient of a postdoctoral fellowship (code SPF20170938626) from the Fondation pour la Recherche Médicale (FRM). Further support was through a European Research Council Consolidator grant: CoG EVO-MEIO 681946 to K.B.

## Contributions

P.B. designed the study with contributions from L.Q., K.B., and V.C. P.B. analyzed the whole genome resequencing data with help from L.Q. B.H. obtained and sequenced the fosmid clones. P.B. performed the assembly of the fosmid sequences as well as the identification and characterization of the *ATCOPIA78* insertion. P.B. and V.C. wrote the manuscript. All authors revised the manuscript.

## Corresponding authors

Correspondence to Vincent Colot.

